# Activated kinesin-1 assembles into a dimer-of-dimers

**DOI:** 10.64898/2025.12.23.696152

**Authors:** Kyoko Chiba

## Abstract

Kinesin-1 mediates intracellular cargo transport along microtubules. The canonical kinesin-1 complex is composed of two KIF5 motor subunits (also known as kinesin heavy chain; KHCs) and two cargo-binding kinesin light chains (KLCs), and KIF5 is generally thought to function as a homodimer. Whether kinesin-1 forms higher-order assemblies beyond the dimer has remained unclear. Here, we show that the neuronal isotype KIF5C assembles into tetramers through reversible association of two dimers. Gel filtration and mass photometry demonstrate that KIF5C partitions between dimeric and tetrameric forms *in vitro*. Deletion of the autoinhibitory elbow region strongly shifts this distribution toward tetramers, indicating that conformational activation promotes intermolecular assembly. Tetramer formation requires the C-terminal tail but remains compatible with KLC binding. Single-molecule motility assays reveal that tetramer-enriched KIF5C exhibits increased landing rates and longer run lengths compared with the dimeric form. These findings identify reversible tetramerization as a previously unrecognized property of kinesin-1 and suggest that conformational activation not only relieves intramolecular inhibition but also promotes higher-order assembly, revealing an additional layer of regulation that may tune motor activity in neurons.

## Introduction

Kinesin-1 motors (KIF5 family proteins) are plus-end directed microtubule motors that play key roles in various intracellular events including organelle positioning, microtubule organization, and cytoplasmic streaming (Hirokawa et al., 2009, Yan et al., 2013, Lu et al., 2016). The canonical kinesin-1 complex is a heterotetramer composed of two kinesin heavy chains (KHCs; also known as KIF5) and two kinesin light chains (KLCs) (Bloom et al., 1988, Kuznetsov et al., 1988, Johnson et al., 1990). Each KIF5 polypeptide contains an N-terminal motor domain, a stalk domain formed by sequential coiled-coil domains, and a C-terminal tail (Hirokawa et al., 1989, de Cuevas et al., 1992). In addition to the KIF5-KLC complex, KIF5 can exist as homodimers lacking KLCs (Hackney et al., 1991, DeLuca et al., 2001).

Oligomerization regulates several motor proteins. Members of the kinesin superfamily can transition between monomeric and dimeric states, markedly enhancing motility (Soppina et al., 2014, Ren et al., 2016, Kita et al., 2023, Fan and McKenney, 2023, Niwa et al., 2024a, Niwa et al., 2024b). Dysregulation of this transition impairs cargo transport (Chiba et al., 2019), and contributes to disease (Chiba et al., 2023, Niwa et al., 2024b). In cytoplasmic dynein-1 (dynein), recruitment of a second dynein dimer to dynactin-adaptor complexes increases force production and velocity (Grotjahn et al., 2018, Urnavicius et al., 2018, Elshenawy et al., 2019, Chaaban and Carter, 2022). These examples illustrate how regulated changes in oligomeric state modulate motor output.

Kinesin-1 is well known to be regulated through autoinhibition (Hackney et al., 1992, Verhey et al., 1998). This inhibition is mediated by intramolecular interactions which promote folding of the molecule in half, preventing ATP hydrolysis and motility (Stock et al., 1999, Coy et al., 1999, Kaan et al., 2011). In vivo and in vitro experiments have shown that binding of cargo adaptors or microtubule-associated proteins relieves this inhibition (Verhey et al., 2001, Ferro et al., 2022, Canty et al., 2023, Chiba et al., 2022, Blasius et al., 2007, Sun et al., 2011, Barlan et al., 2013, Fu and Holzbaur, 2013, Twelvetrees et al., 2019, Henrichs et al., 2020, Hooikaas et al., 2019). Recently, a flexible linker located within the stalk region of the KHC termed the "elbow" was proposed to act as the hinge that allows the molecule to fold into the autoinhibited conformation (Weijman et al., 2022, Tan et al., 2023). Although deletion of the elbow induces molecular extension and motor activation in cells, its effects on oligomeric state have not been examined biophysically.

Mammals express three KIF5 isotypes, KIF5A, KIF5B and KIF5C, which share highly conserved motor and coiled-coil domains but differ in their C-terminal regions. KIF5A and KIF5C are expressed in nervous tissue, whereas KIF5B is ubiquitous (Aizawa et al., 1992, Nakagawa et al., 1997, Kanai et al., 2000). In previous work comparing the motility of these KIF5 isotypes, we noted the presence of higher-order KIF5 oligomers during protein preparation (Chiba et al., 2022). Oligomeric species, including tetramers, were not completely excluded and were present in subsequent analyses (Fig. 1I and Fig. S2 of Chiba et al) (Chiba et al., 2022). Their consistent appearance suggested a tendency of KIF5 to form higher-order assemblies. Recent observations from another research group have further raised the possibility that KIF5 molecules can form tetrameric assemblies (Carrington et al., 2024). However, the molecular basis, regulation and functional relevance of such oligomers remains unknown. In this study, we examine the oligomeric state of the neuronal isotype KIF5C and show that KIF5C can form tetramers through association of two dimers and that conformational activation strongly promotes this intermolecular assembly.

**Figure 1.**
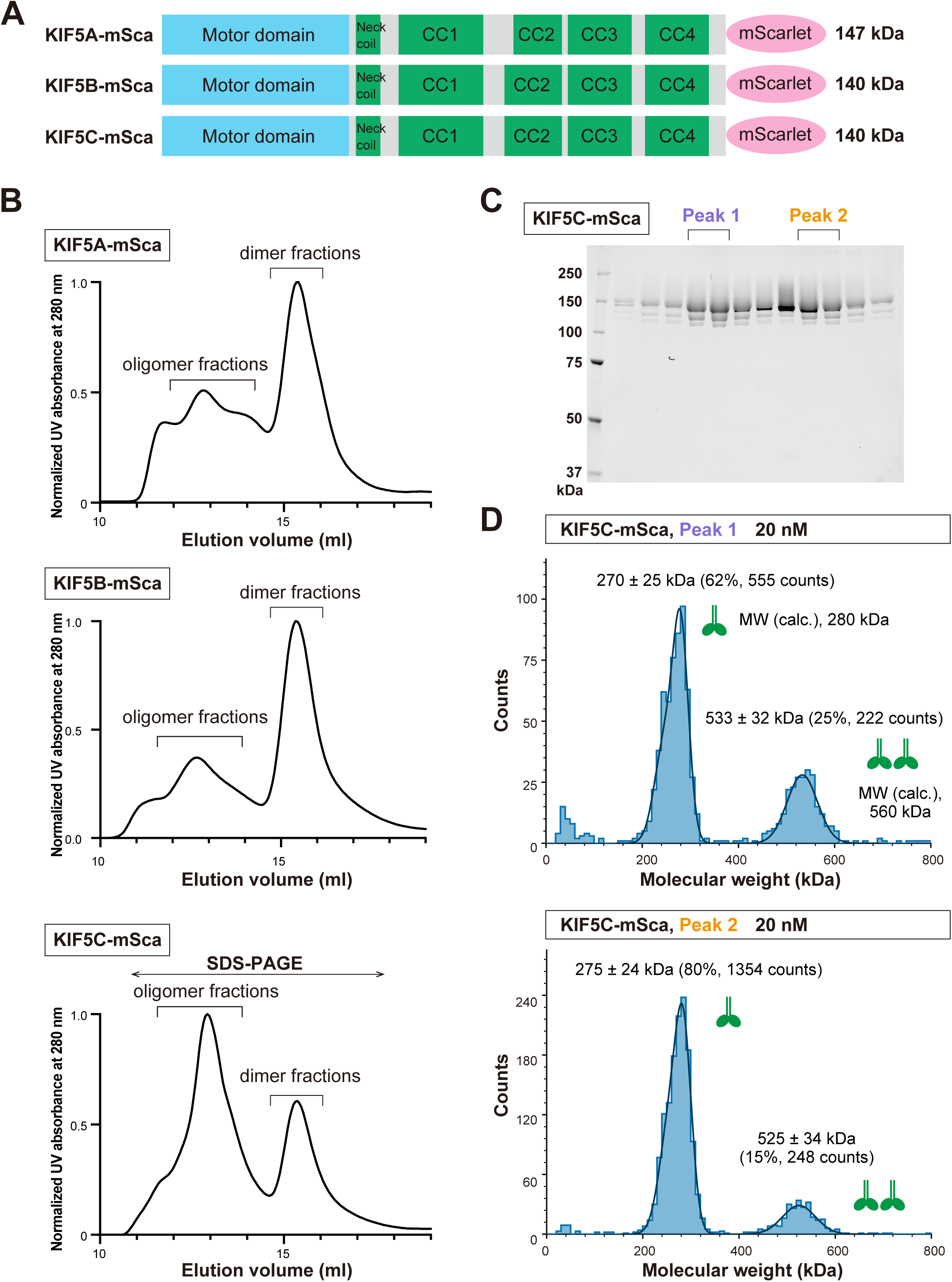
A subset of KIF5C assembles into a tetrameric state *in vitro*. **(A)** Schematic representation of human KIF5 isotypes fused with C-terminal mScarlet and Strep-tag II (KIF5A-mSca, KIF5B-mSca and KIF5C-mSca). Calculated molecular weights including mScarlet and Strep-tag II are shown. CC, coiled-coil domain. **(B)** Gel filtration chromatography profile of KIF5-mSca proteins. Fractions corresponding to dimeric and oligomeric fractions are marked by inverted U-shapes. KIF5C-mSca fractions indicated by a double-headed arrow were analyzed by SDS–PAGE. **(C)** SDS-PAGE of the KIF5C-mSca fractions from (B). Peak 1 (purple) and Peak 2 (orange) are marked by inverted U-shapes. Molecular weights (kDa) are shown on the left. **(D)** Mass photometry of KIF5C-mSca Peak 1 and Peak 2 at 20 nM. Histograms show molecular-weight distributions with Gaussian fits (mean ± SD). Particle counts and relative percentages are indicated. Green diagrams represent KIF5C molecules corresponding to the inferred oligomeric states. Calculated molecular weights are shown.

## Results

### Oligomerization property of KIF5 isotypes

To examine whether oligomerization is a general property of the KIF5 family, we first analyzed the oligomeric states of all three mammalian KIF5 isotypes (KIF5A, KIF5B, and KIF5C) using recombinant full-length proteins expressed in Sf9 cells (Fig. 1A) (Chiba et al., 2022). Each isotype was purified by affinity chromatography followed by gel filtration at micromolar concentrations. Gel-filtration analysis revealed that all three KIF5 isotypes eluted as multiple peaks, indicating oligomeric heterogeneity *in vitro* (Fig. 1B). In each case, a distinct peak was observed around an elution volume of ∼15.5 mL, which corresponds to the dimeric species as previously determined by SEC-MALS analysis (Chiba et al., 2022). In addition to this major dimer peak, earlier-eluting peaks were consistently detected for all isotypes, suggesting the presence of higher-molecular-weight species. Among the three isotypes, KIF5C exhibited the most pronounced oligomerization behavior. KIF5C-mSca was separated into two major gel-filtration peaks at approximately 13 mL and 15.5 mL with comparable UV absorbance (Fig. 1B and C). Because of this clear separation and reproducible elution profile, we focused subsequent analyses on KIF5C to define the molecular basis, regulation, and functional consequences of its assembly.

### KIF5C forms dimers and tetramers

To define the oligomeric states of KIF5C, gel-filtration peak fractions of KIF5C-mSca were analyzed by mass photometry (Fig. 1D). At 20 nM, particles from the earlier peak (Peak 1) resolved into two major populations corresponding to a dimer (270 ± 25 kDa, mean ± SD) and a tetramer (533 ± 32 kDa, mean ± SD), accounting for 62% and 25% of particles, respectively (Fig. 1D). The later peak (Peak 2) also contained both dimeric (275 ± 24 kDa, mean ± SD) and tetrameric species (525 ± 34 kDa, mean ± SD), comprising 80% and 15% of total particles (Fig. 1D). These results demonstrate that KIF5C exists in both dimeric and tetrameric forms. The presence of both species within each fraction suggests reversible exchange between the two forms.

### KIF5C dimers associate dynamically to form tetramers

To test whether the dimeric and tetrameric species interconvert, we performed sequential gel filtration. Each peak fraction from the initial gel filtration was stored at 4°C for 48 h and then re-analyzed under identical conditions (Fig. 2A). In the secondary gel filtration, both initial Peak 1 and Peak 2 yielded indistinguishable chromatograms. Re-injection of either fraction again produced two peaks corresponding to the original pattern (Fig. 2B). These findings indicate reversible interconversion between dimeric and tetrameric forms, consistent with tetramers arising from association of two dimers.

**Figure 2.**
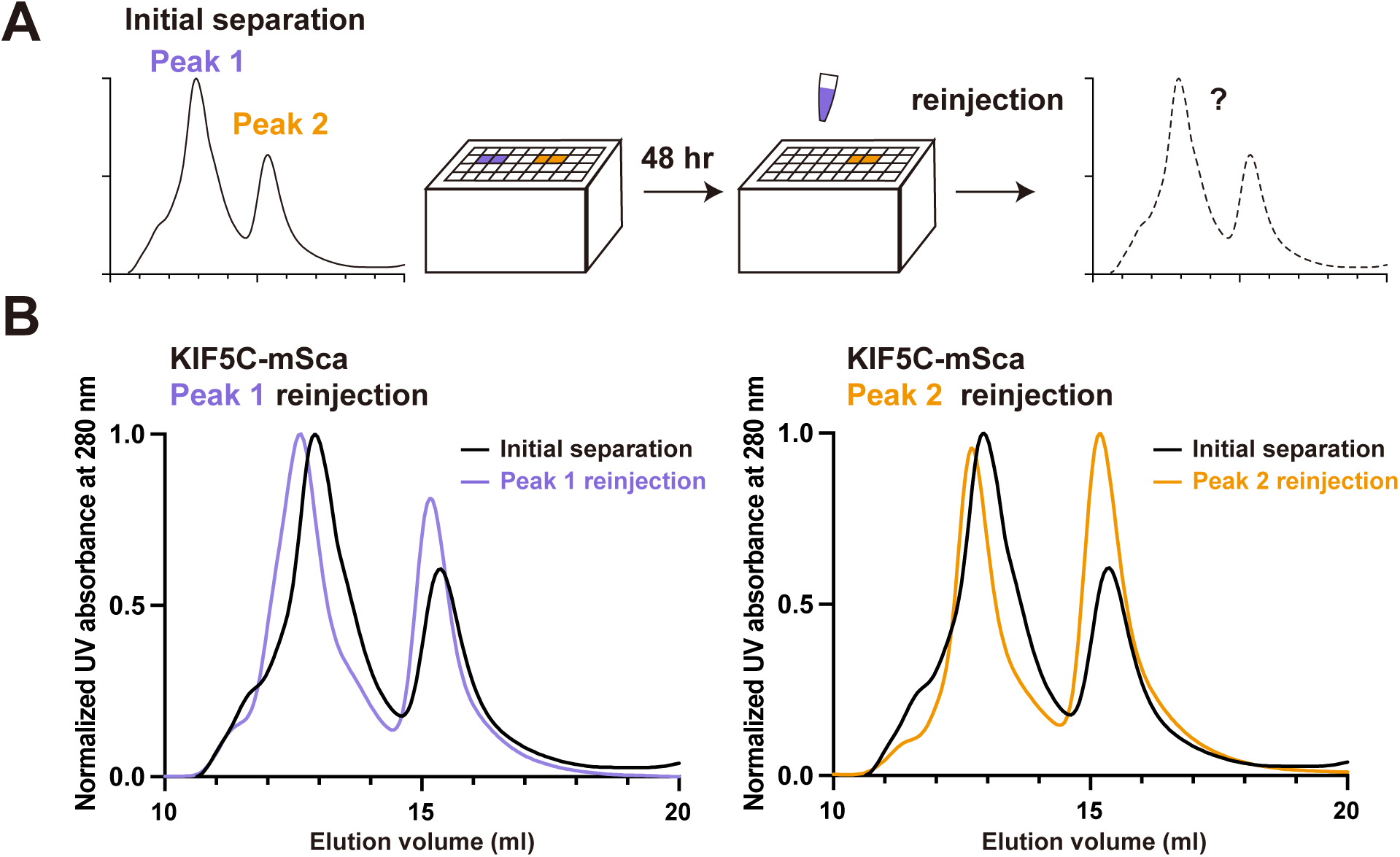
KIF5C dimers associate dynamically to form tetramers. **(A)** Schematic representation of the experimental procedure. KIF5C-mSca proteins were first separated by gel filtration chromatography (initial separation). Eluted fractions were stored for 48 h at 4°C and then re-injected for secondary gel filtration analysis. **(B)** Gel filtration profiles of KIF5C-mSca. Black lines represent the initial separation, and purple and orange lines represent the secondary separation of Peak 1 and Peak 2, respectively.

### Elbow deletion enhances tetramer formation of KIF5C

KIF5 proteins are regulated by autoinhibition, which involves folding at a flexible linker region termed the "elbow" (Weijman et al., 2022). Deletion of this region (Δelbow) has been shown to promote an extended molecular conformation (Weijman et al., 2022). To test how conformational activation affects the oligomeric state, we generated a Δelbow mutant (KIF5C(Δelbow)-mSca, 137 kDa) by deleting residues from Glu675 to Arg692, corresponding to the region removed in the original report (Fig. 3A). Purified KIF5C(Δelbow)-mSca eluted as a single peak corresponding to Peak 1 of wild-type KIF5C (Fig. 3B, 3C), consistent with previous findings (Weijman et al., 2022). Although previous work concluded that the Δelbow mutant is dimeric, mass photometry revealed that KIF5C(Δelbow)-mSca predominantly assembles into tetramers. Mass photometry detected a species consistent with a tetramer (547 ± 49 kDa, mean ± SD) (Fig. 3D, 3I). These results indicate that release from autoinhibition markedly enhances assembly of KIF5C dimers into tetramers.

**Figure 3.**
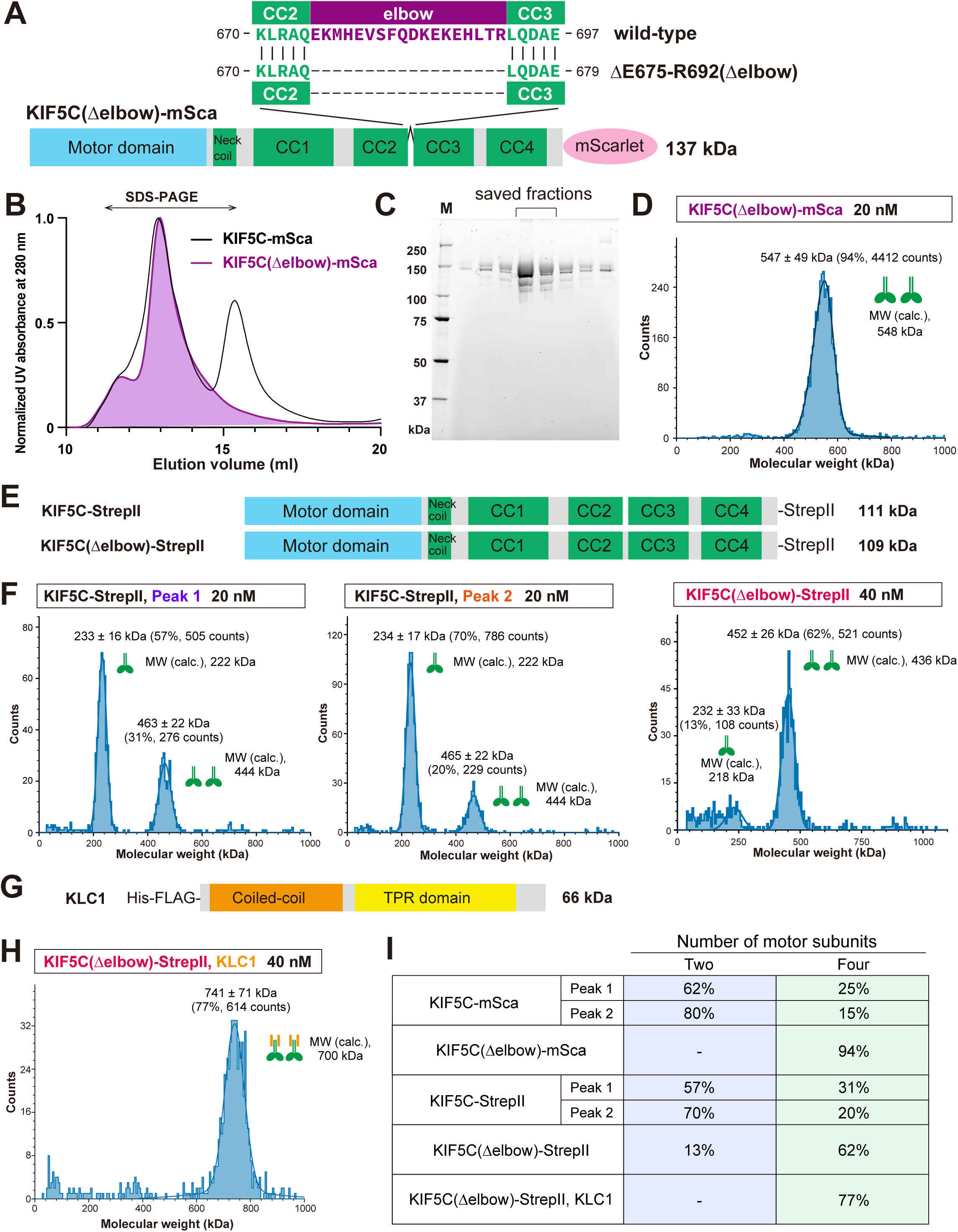
Elbow deletion promotes tetramer formation of KIF5C. **(A)** Schematic representation of the Δelbow mutant of KIF5C, in which residues from E675 to R692 are deleted. The calculated molecular weight including mScarlet and Strep-tag II is shown. CC, coiled-coil domain. **(B)** Gel filtration chromatography profile of KIF5C(Δelbow)-mSca. Black line represents wild-type KIF5C (KIF5C-mSca) replotted from Fig. 1B. Fractions indicated by a double-headed arrow were analyzed by SDS–PAGE. **(C)** SDS-PAGE of fractions from (B). The elution peak is indicated by an inverted U-shape. Molecular weight markers (kDa) are shown on the left. **(D)** Mass photometry of KIF5C(Δelbow)-mSca at 20 nM. Histogram shows molecular weight distributions with a Gaussian fit (mean ± SD). Particle counts and relative percentages are indicated. Green diagrams represent KIF5C assemblies corresponding to the inferred oligomeric states. Calculated molecular weight is shown. **(E)** Schematic representation of KIF5C-StrepII and KIF5C(Δelbow)-StrepII. Calculated molecular weights including tags are shown on the right. CC, coiled-coil domain. **(F)** Mass photometry of KIF5C-StrepII (20 nM) and KIF5C(Δelbow)-StrepII (40 nM). Histograms show molecular weight distributions with Gaussian fits (mean ± SD). Particle counts and relative percentages are indicated. Green diagrams represent KIF5C molecules corresponding to the inferred oligomeric states. Calculated molecular weights for the inferred oligomeric states are shown. **(G)** Schematic representation of KLC1. Calculated molecular weight including tags is shown. CC, coiled-coil domain. TPR domain, tetratricopeptide repeat domain. **(H)** Mass photometry of KIF5C(Δelbow)-StrepII, KLC1 at 40 nM. Histograms show molecular weight distributions with a Gaussian fit (mean ± SD). Particle counts and relative percentages are indicated. Green diagrams represent KIF5C and KLC1 molecules corresponding to the inferred oligomeric states. **(I)** Table summarizing the oligomeric states of KIF5C. Relative percentages of each oligomeric state corresponding to two or four motor subunits are shown.

### Removal of the fluorescent tag does not affect KIF5C tetrameric assembly

Because fluorescent proteins can sometimes promote artificial multimerization, we examined whether the mScarlet tag influenced KIF5C tetrameric assembly. The untagged construct (KIF5C-StrepII, 111 kDa) was expressed in insect cells and purified (Fig. 3E, S1A, S1B). Similar to KIF5C-mSca, KIF5C-StrepII eluted as two peaks on gel filtration (Fig. S1A, S1B). Mass photometry of Peak 1 revealed tetrameric (463 ± 22 kDa) and dimeric (233 ± 16 kDa) species, accounting for 57% and 31% of total particles (Fig. 3F, 3I). Peak 2 contained 70% dimers (234 ± 17 kDa) and 20% tetramers (465 ± 22 kDa) (Fig. 3F, 3I). These results indicate that C-terminal fluorescent tags do not alter KIF5C tetramer formation. Likewise, the non-fluorescent KIF5C Δelbow mutant (KIF5C(Δelbow)-StrepII, 109 kDa) eluted as a single peak corresponding to Peak 1 of KIF5C-StrepII (Fig. S1A, S1B). Mass photometry of KIF5C(Δelbow)-StrepII detected 13% dimers (232 ± 33 kDa, mean ± SD) and 62% tetramers (452 ± 26 kDa, mean ± SD) (Fig. 3F and 3I). These findings further support that KIF5C forms tetramers through association of two dimers, rather than through artifacts caused by fluorescent tagging.

### Tetramer formation is compatible with KLC binding

KIF5 proteins form heterotetrameric kinesin-1 complexes with KLCs. KLCs interact with both coiled-coil and flexible regions of KIF5 and generally inhibit motor activity. To determine whether KLC binding affects KIF5C intermolecular association, we co-expressed KIF5C(Δelbow)-StrepII with His-FLAG-KLC1 (66 kDa) and purified the complex by tandem affinity purification (Fig. 3G and S2A-B). The KIF5C(Δelbow)-KLC1 complex eluted as a single peak similar to KIF5C(Δelbow)-StrepII alone (Fig. S2A). SDS-PAGE confirmed the presence of both KIF5C and KLC1 in the peak (Fig. S2B). Mass photometry revealed an average molecular weight of 741 ± 71 kDa, consistent with an assembly containing four copies of both KIF5C and KLC polypeptides (Fig. 3H and 3I). These results indicate that KLC binding does not disrupt KIF5C self-association and that tetrameric KIF5C retains its ability to bind KLC.

### Tetrameric KIF5C shows increased motility on microtubules

To assess how tetramer formation influences KIF5C motility, single-molecule assays were performed using total internal reflection fluorescence (TIRF) microscopy. Two gel-filtration peaks of wild-type KIF5C and the Δelbow mutant were compared. At 20 nM in the presence of 2 mM ATP, Peak 2 of wild-type KIF5C showed few binding events on microtubules, consistent with strong autoinhibition (Chiba et al., 2022). In contrast, Peak 1 exhibited more frequent binding (Fig. 4A, E). Their mean landing rates were 0.011 and 0.0047 molecules/μm/s for Peak 1 and 2, respectively (Fig. 4E). Particles in Peak 1 moved for longer durations and distances than those in Peak 2, although their velocities were not significantly different (Fig. 4G). The Δelbow mutant (KIF5C(Δelbow)-mSca) displayed markedly increased binding to microtubules at 20 nM, the same concentration used for the mass photometry (Fig. 4B). Motors frequently accumulated at microtubule ends, and prolonged observation (∼ 10 min) revealed clusters of motors that remained motile along microtubules (Fig. 4C). Because individual motors could not be distinguished at this concentration, assays were repeated at 2 nM (Fig. 4D). When normalized to motor concentration, the Δelbow mutant showed a significantly higher landing rate than the wild-type (Fig. 4F). The mutant frequently exhibited movement accompanied by diffusive motion, leading to longer dwell times compared with wild type, whereas the run length was comparable to that of Peak 1 (Fig. 4H). These data indicate that tetramer-enriched KIF5C exhibits enhanced motility relative to the dimeric form.

**Figure 4.**
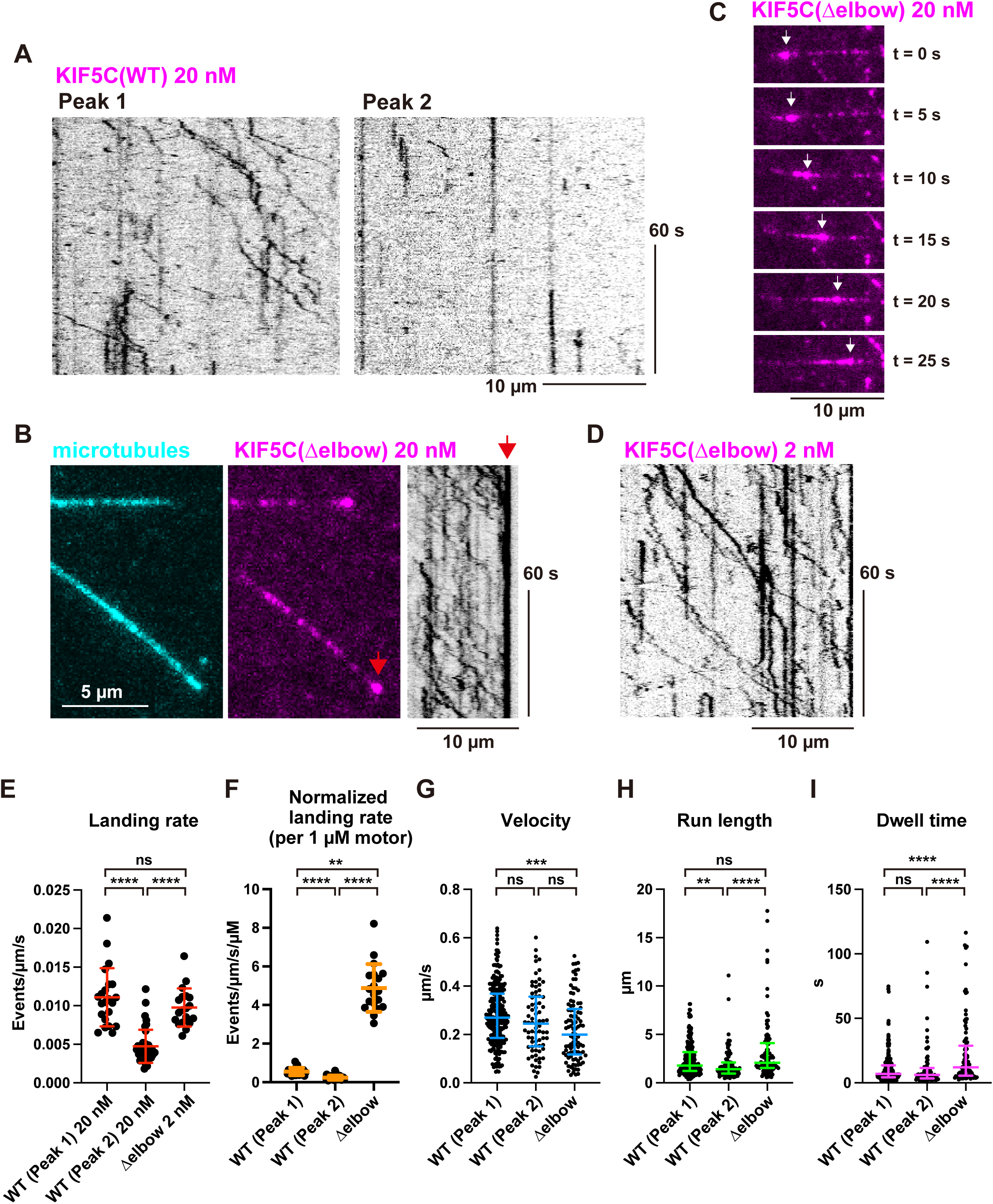
Tetrameric KIF5C exhibits enhanced motility on microtubules. **(A)** Representative kymographs showing the motility of KIF5C-mSca (Peak 1) and (Peak 2) at 20 nM in the presence of 2 mM ATP. Images were acquired at 2 frames per second (fps). Horizontal and vertical scale bars represent 10 µm and 60 s, respectively. **(B)** Representative images and a kymograph showing KIF5C(Δelbow)-mSca at 20 nM in the presence of 2 mM ATP. Cyan, microtubules; magenta, KIF5C(Δelbow)-mSca. Red arrows indicate microtubule tips. Images were acquired at 2 fps within 5 min after loading the motor solution into the chamber. Horizontal and vertical scale bars represent 10 µm and 60 s, respectively. **(C)** Representative time-lapse images showing a KIF5C(Δelbow)-mSca aggregate in the same chamber as in (B). Images were acquired 10 min after loading the motor solution. White arrows indicate an aggregate moving along a microtubule. Images are shown every 5 s. Scale bar, 10 µm. **(D)** Representative kymographs showing the motility of KIF5C(Δelbow)-mSca at 2 nM in the presence of 2 mM ATP. Images were acquired at 2 fps. Horizontal and vertical scale bars represent 10 µm and 60 s, respectively. **(E)** Dot plots showing the landing rate of KIF5C-mSca (Peak 1), KIF5C-mSca (Peak 2) and KIF5C(Δelbow)-mSca. The number of molecules that bound to microtubules was normalized to microtubule length (µm) and time (s). All motile, immotile and non-directional molecules were counted. Each dot represents one data point. Bars represent mean ± SD. n = 22, 40 and 19 microtubules, respectively. Kruskal–Wallis test followed by Dunn’s multiple-comparison test. ****, p < 0.0001; ns, not statistically significant. **(F)** Dot plots showing the landing rate normalized by motor concentration (µM). The number of molecules bound to microtubules was normalized to microtubule length (µm), time (s) and motor concentration (µM). Bars represent mean ± SD. n = 22, 40 and 19 microtubules, respectively. Kruskal– Wallis test followed by Dunn’s multiple-comparison test. **, p < 0.01; ****, p < 0.0001; ns, not statistically significant. **(G)** Dot plots showing the velocity of KIF5C-mSca (Peak 1), KIF5C-mSca (Peak 2) and KIF5C(Δelbow)-mSca. Each dot represents one data point. Bars represent mean ± SD. n = 258, 77 and 97, respectively. Kruskal–Wallis test followed by Dunn’s multiple-comparison test. ***, p < 0.001; ns, not statistically significant. **(H)** Dot plots showing the run length of KIF5C-mSca (Peak 1), KIF5C-mSca (Peak 2) and KIF5C(Δelbow)-mSca. Each dot represents one data point. Bars represent median value and interquartile range. n = 258, 77 and 97, respectively. Kruskal–Wallis test followed by Dunn’s multiple-comparison test. **, p < 0.01; ****, p < 0.0001; ns, not statistically significant. **(I)** Dot plots showing the dwell time of KIF5C-mSca (Peak 1), KIF5C-mSca (Peak 2) and KIF5C(Δelbow)-mSca. Each dot represents one data point. Bars represent median value and interquartile range. n = 258, 77 and 97, respectively. Kruskal–Wallis test followed by Dunn’s multiple-comparison test. ****, p < 0.0001; ns, not statistically significant.

### The C-terminal tail is required for KIF5C assembly

Truncated KIF5 constructs containing only the motor domain and part of the stalk domain (e.g., K420 or K560) are widely used in motility assays, however, tetrameric forms have not been observed for these fragments (Vale et al., 1996, Friedman and Vale, 1999, Hooikaas et al., 2019, Chiba et al., 2022, Shima et al., 2018). We therefore hypothesized that the C-terminal region, which is absent from these constructs, contributes to KIF5C tetramer formation. To identify the region responsible for dimer-dimer association, we deleted the C-terminal 43 amino acids (Fig. 5A). C-terminally truncated versions of wild-type and Δelbow mutants, KIF5C(Δtail)-mSca (135 kDa) and KIF5C(ΔtailΔelbow)-mSca (133 kDa), were expressed and purified. KIF5C(Δtail)-mSca showed two UV peaks on gel filtration (Fig. 5B), but SDS-PAGE revealed that the first UV peak contained little protein relative to absorbance, indicating that most protein eluted in the second peak. Proteins eluting near the void volume were excluded from analysis. Mass photometry of the second peak showed that KIF5C(Δtail)-mSca was predominantly dimeric (281 ± 24 kDa, mean ± SD; 93% of total particles) (Fig. 5C). Next, we examined whether C-terminal deletion affected the Δelbow mutant. The gel-filtration profile of KIF5C(ΔtailΔelbow)-mSca was similar to that of KIF5C(Δelbow) (Fig. 5D). However, mass photometry revealed that KIF5C(ΔtailΔelbow) primarily existed as dimers (251 ± 36 kDa, mean ± SD; 91% of total particles) (Fig. 5E). These results demonstrate that the C-terminal tail is required for tetramer formation.

**Figure 5.**
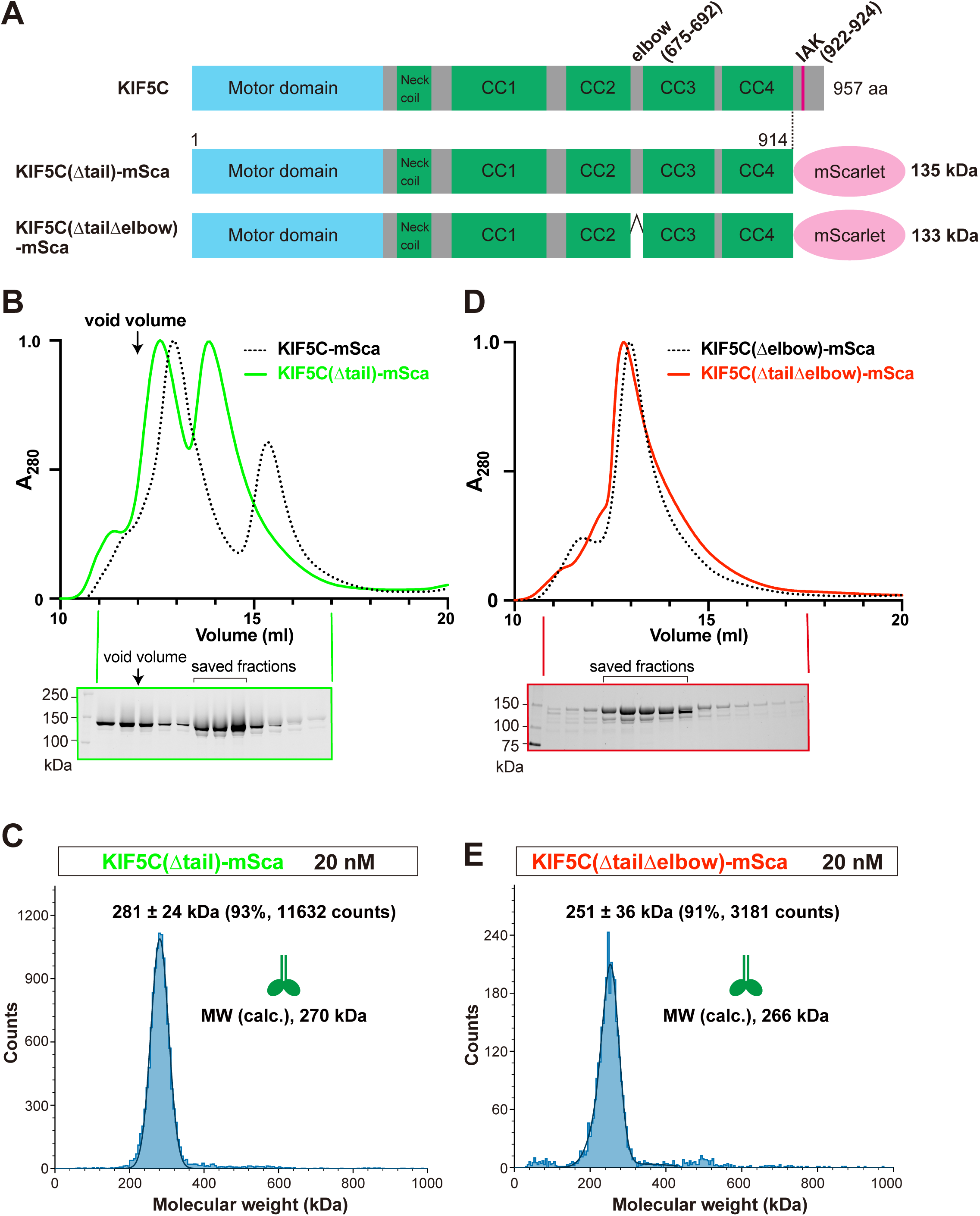
The C-terminal tail is required for KIF5C tetramerization. **(A)** Schematic representation of KIF5C(Δtail)-mSca and KIF5C(ΔtailΔelbow)-mSca. The elbow region and IAK motif are indicated together with the corresponding amino-acid numbers. Calculated molecular weights including mScarlet and Strep-tag II are shown. CC, coiled-coil domain. **(B)** Gel filtration chromatogram of KIF5C(Δtail)-mSca. Dotted line represents KIF5C-mSca replotted from Fig. 1B. Arrow indicates the void volume of the column (12 mL). SDS–PAGE of the eluted fractions is shown below. Molecular weight markers (kDa) are shown on the left. Saved fractions are marked by an inverted U-shape. **(C)** Gel filtration chromatogram of KIF5C(ΔtailΔelbow)-mSca. Dotted line represents KIF5C(Δelbow)-mSca replotted from Fig. 3B. SDS–PAGE of the eluted fractions is shown below. Molecular weight markers (kDa) are shown on the left. Saved fractions are marked by an inverted U-shape. **(D, E)** Mass photometry of KIF5C(Δtail)-mSca (D) and KIF5C(ΔtailΔelbow)-mSca (E) at 20 nM. Histograms show molecular weight distributions with Gaussian fits (mean ± SD). Particle counts and relative percentages are indicated. Green diagrams represent KIF5C molecules corresponding to the inferred oligomeric states. Calculated molecular weights are shown.

**Figure 6.**
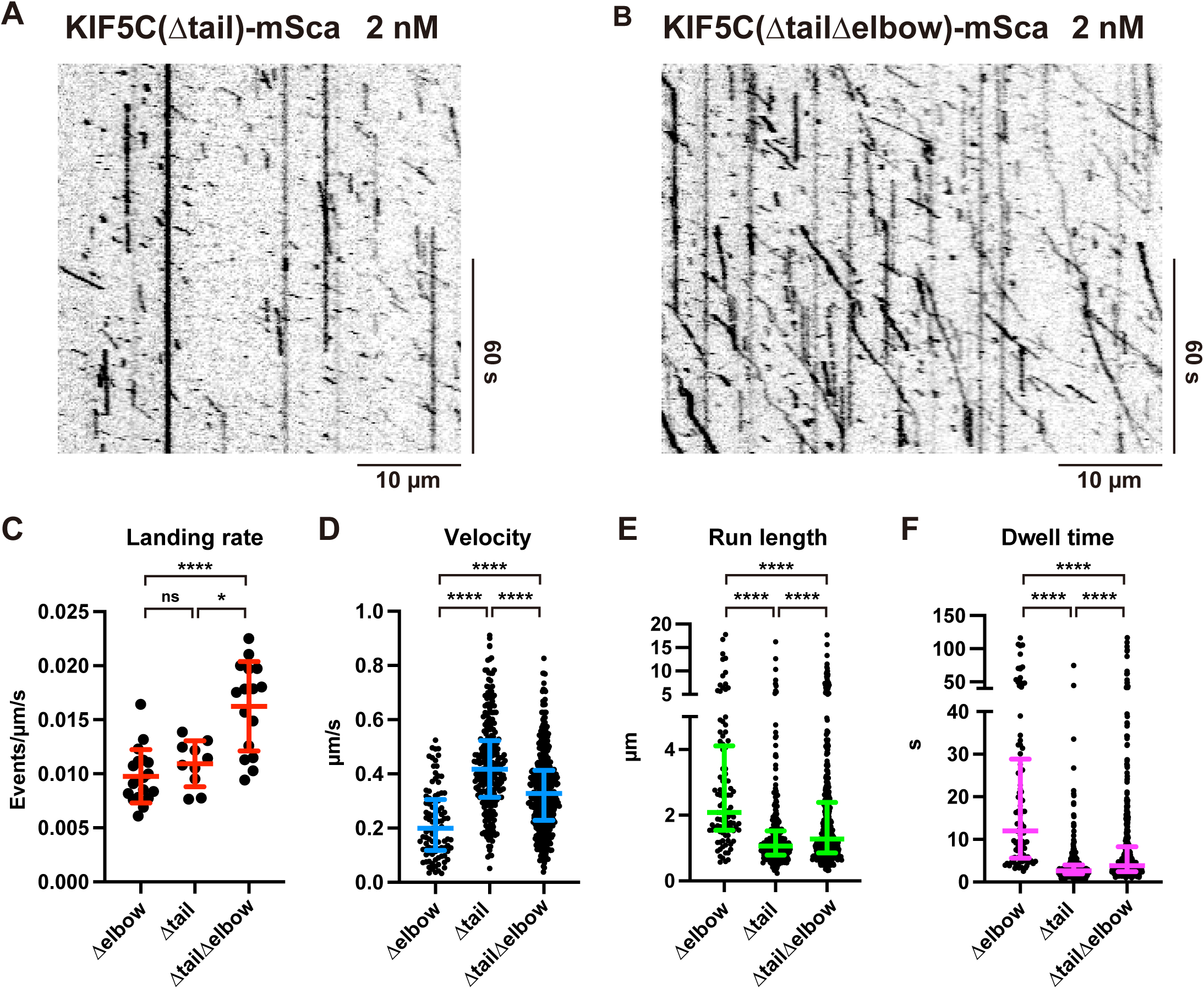
C-terminal deletion alters the motility of elbow-deleted KIF5C. **(A)** Representative kymographs showing the motility of KIF5C(Δtail)-mSca at 2 nM in the presence of 2 mM ATP. Images were acquired at 2 fps. Horizontal and vertical scale bars represent 10 µm and 60 s, respectively. **(B)** Representative kymographs showing the motility of KIF5C(ΔtailΔelbow)-mSca at 2 nM in the presence of 2 mM ATP. Images were acquired at 2 fps. Horizontal and vertical scale bars represent 10 µm and 60 s, respectively. **(C)** Dot plots showing the landing rate of KIF5C(Δtail)-mSca and KIF5C(ΔtailΔelbow)-mSca. The number of molecules that bound to microtubules was normalized to microtubule length (µm) and time (s). All motile, immotile and non-directional molecules were counted. Data for KIF5C(Δelbow)-mSca were replotted from Fig. 4E. Each dot represents one data point. Bars represent mean ± SD. n = 19, 10 and 16 microtubules, respectively. Kruskal–Wallis test followed by Dunn’s multiple-comparison test. *, p < 0.05; ****, p < 0.0001; ns, not statistically significant. **(D)** Dot plots showing the velocity of KIF5C(Δtail)-mSca and KIF5C(ΔtailΔelbow)-mSca. Each dot represents one data point. Bars represent mean ± SD. n = 97, 275 and 442, respectively. Kruskal–Wallis test followed by Dunn’s multiple-comparison test. ****, p < 0.0001. **(E)** Dot plots showing the run length of KIF5C(Δtail)-mSca and KIF5C(ΔtailΔelbow)-mSca. Each dot represents one data point. Bars represent median value and interquartile range. n = 97, 275 and 442, respectively. Kruskal–Wallis test followed by Dunn’s multiple-comparison test. ****, p < 0.0001. **(F)** Dot plots showing the dwell time of KIF5C(Δtail)-mSca and KIF5C(ΔtailΔelbow)-mSca. Each dot represents one data point. Bars represent median value and interquartile range. n = 97, 275 and 442, respectively. Kruskal–Wallis test followed by Dunn’s multiple-comparison test. ****, p < 0.0001.

### C-terminal deletion alters the motility of activated KIF5C

Using KIF5C(Δtail) and KIF5C(ΔtailΔelbow), which lack detectable tetramers, we next examined how the C-terminus-mediated assembly influences KIF5C motility. In single-molecule assays at 2 nM, KIF5C(Δtail) displayed a landing rate comparable to the Δelbow mutant but exhibited shorter and faster movements. Diffusive motion observed in the Δelbow mutant was absent in KIF5C(Δtail). Similarly, KIF5C(ΔtailΔelbow) showed no diffusive movement. Unlike KIF5C(Δtail), KIF5C(ΔtailΔelbow) contained a subset of slower molecules that often moved longer distances, resulting in a lower mean velocity. The landing rate of KIF5C(ΔtailΔelbow) was significantly higher than that of KIF5C(Δtail), consistent with previous findings that combined mutations in the IAK and elbow regions produce greater activation than either mutation alone (Tan et al., 2023).

## Discussion

### KIF5C forms tetramers through association between dimers

This study demonstrates that KIF5C forms tetramers through reversible association of two dimers. Gel filtration and mass photometry revealed the coexistence of dimeric and tetrameric species and supported interconversion between these forms. This self-association extends previous observations of higher-order assemblies of KIF5 family proteins, including reports of tetrameric KIF5A (Chiba et al., 2022, Carrington et al., 2024). Although the molecular mechanism may differ among KIF5 isotypes, the presence of tetramers in both KIF5A and KIF5C suggests that the capacity for higher-order assembly is an intrinsic property of kinesin-1 motors.

### Conformational activation promotes dimer-dimer association

Deletion of the autoinhibitory elbow region markedly increased tetramer formation, indicating a functional link between conformational activation and oligomeric assembly. Previous studies have shown that elbow deletion induces an extended molecular conformation of kinesin-1. These results suggest that conformational activation not only relieves intramolecular autoinhibition but also biases the equilibrium toward intermolecular assembly.

The requirement of the C-terminal tail for tetramer formation further refines this model. In the autoinhibited state, the tail is thought to be constrained through intramolecular interactions. Conformational extension may expose interaction surfaces within the C-terminal region, enabling contacts between dimers that stabilize the tetrameric state.

Although mass photometry directly demonstrates enhanced tetramer formation upon Δelbow modification, interpretation of the associated gel-filtration peak shift requires caution. The ΔtailΔelbow construct remains dimeric by mass photometry yet exhibits a similar elution shift, consistent with previous reports that conformational extension contributes to changes in hydrodynamic behavior (Weijman et al., 2022). Thus, the elution profile of the Δelbow mutant likely reflects a combination of conformational change and tetramer formation.

### Higher-order assembly as a regulatory strategy for kinesin-1

Regulation of motor activity through higher-order assembly is observed in other cytoskeletal motors. Cytoplasmic dynein, for example, exhibits enhanced force production and velocity when two dynein dimers are recruited into a single transport complex (Grotjahn et al., 2018, Urnavicius et al., 2018, Elshenawy et al., 2019, Chaaban and Carter, 2022). A related phenomenon has also been reported for myosin VI, which undergoes oligomerization upon binding to cargo membranes (Montanarella et al., 2025). Although underlying mechanisms differ, these examples illustrate that assembly of motor dimers can serve as an effective strategy to enhance motility.

By analogy, known activators of kinesin-1, including cargo adaptor proteins and microtubule-associated proteins, may promote release from autoinhibition and thereby facilitate tetramer formation of KIF5C. Tetramer formation may therefore represent an additional regulatory step downstream of conformational activation, modulating the collective behavior of motors once activated.

### Orientation of tetramer assembly

Whether two KIF5C dimers associate in a parallel or antiparallel orientation remains unknown. Although structural information is lacking, several observations favor a parallel association model. Tetramer-enriched KIF5C exhibits increased landing rates, prolonged dwell times, and longer run lengths relative to the dimeric form, consistent with a configuration in which multiple motor domains engage the same microtubule. In contrast, antiparallel association would be expected to promote microtubule crosslinking or sliding, as observed for kinesin-5 motors, Eg5 (Kapitein et al., 2005). Such activity was not detected for the KIF5C Δelbow mutant under the conditions tested (data not shown). We therefore favor a parallel association model. Definitive resolution, however, will require structural analysis.

### Possible role of post-translational modifications

Tetramer formation was readily observed for insect cell-expressed KIF5C but was not detected in previous studies using bacterially expressed KIF5C(Δelbow) (Weijman et al., 2022). This discrepancy raises the possibility that eukaryotic post-translational modifications, such as phosphorylation, may facilitate or stabilize inter-dimer interactions. Identification of such modifications and elucidation of their regulatory roles will be an important direction for future studies.

### Physiological relevance and implications for kinesin-1 regulation

The physiological relevance of KIF5C tetramers in cells remains to be established. Early biochemical studies of native kinesin-1, which predominantly relied on motors purified from nervous tissue, did not report higher-order assemblies (Vale et al., 1985, Kuznetsov and Gelfand, 1986, Kuznetsov et al., 1988, Bloom et al., 1988, Hirokawa et al., 1989, Hackney et al., 1991). This absence may partly reflect the predominance of KIF5B, which is the most abundant kinesin-1 isotype in nervous tissue (Kanai et al., 2000), as well as the use of soluble extracts lacking motors associated with cargo. Our *in vitro* analyses indicate that KIF5B exhibits a weaker tendency toward oligomerization than KIF5C, potentially limiting detection of tetrameric species in earlier preparations.

Because conformational activation strongly promotes tetramer formation *in vitro* (Fig. 3), it is plausible that KIF5C tetramers arise transiently at cargo surfaces where kinesin-1 is locally activated. Punctate cytoplasmic localization of KIF5 observed in cells may reflect localized assembly of active motor complexes (Niwa and Chiba, 2023). Given that aberrant oligomerization of KIF5 has been implicated in neurodegenerative diseases (Pant et al., 2022, Nakano et al., 2022, Chiba and Niwa, 2024), tight regulation of the balance between dimeric and higher-order assemblies is likely critical for neuronal homeostasis.

Together, our findings expand the prevailing model of kinesin-1 regulation. In addition to intramolecular autoinhibition and cargo-mediated activation, kinesin-1 may also be regulated through reversible self-association of motor dimers into higher-order assemblies. In this framework, conformational activation not only releases inhibitory folding but also promotes intermolecular interactions that enhance motor engagement with microtubules. Although the existence and regulation of KIF5C tetramers *in vivo* remain to be established, our study introduces higher-order assembly as an additional dimension of kinesin-1 regulation and provides a conceptual basis for understanding how motor activity may be tuned spatially and temporally in neurons.

### Experimental procedures

#### Plasmids

pACEBac1 human KIF5C-mScarlet-Strep-tag II and pIDS His-FLAG-human KLC1 were described previously. KIF5C(Δelbow)-mScarlet-Strep-tag II, KIF5C-Strep-tag II, KIF5C(Δelbow)-Strep-tag II were generated by PCR and Gibson assembly and cloned into pACEBac1. Primers used for amplifications are listed below. pACEBac1_KIF5C(Δelbow)-StrepII was assembled with pIDS_His-FLAG-HsKLC1 using Cre recombination. The sequence of the plasmids was verified by Sanger sequencing. Plasmids used in the study are summarized in ***Supplementary Table S1*.**

KIF5C(Δ675-692(Δelbow))_F, gagctggcaaagctccgagcccagttgcaggatgctgaagaaatgaagaaggcgctg
KIF5C(Δ675-692(Δelbow))_R, cagcgccttcttcatttcttcagcatcctgcaactgggctcggagctttgccagctc
KIF5C(1-914(Δtail))_F, tgcgggccaagaacatggccggcagtggatcgggtcttga
KIF5C(1-914 (Δtail))_R, tcaagacccgatccactgccggccatgttcttggcccgca
KIF5C-Strep-tag II_F, tggagtcatccacaatttgagaag
KIF5C-Strep-tag II_R, caaattgtggatgactccactcgagtttctggtagtgagtggaatttgaagagctg

#### Expression of recombinant proteins in Sf9 cells

Sf9 cells (Thermo Fisher Scientific) were maintained in Sf900^TM^ II SFM (Thermo Fisher Scientific) at 27°C. DH10Bac cells (Thermo Fisher Scientific) were transformed with the plasmids to generate bacmids. Baculoviruses were produced by transfecting 1 × 10^6^ of Sf9 cells in a 35 mm dish with 10 µg of bacmid using 5 μL of TransIT-Insect (Takara Bio Inc.). Five days post-transfection, the culture supernatant (P1) was collected by centrifugation (15,000 × g, 1 min). For protein expression, 250 mL of Sf9 cells (2 × 10^6^ cells/mL) were infected with 200 µL of P1 virus and cultured for 65 hours at 27°C. Cells were harvested by centrifugation (2,000 × g, 10 min) and stored at −80°C.

#### Purification of recombinant proteins

Cells were suspended in 25 mL of lysis buffer (50 mM HEPES-KOH, 150 mM KCH3COO, 2 mM MgSO4, 10% glycerol, pH 7.5) supplemented with 0.1 mM ATP and protease inhibitors (1 mM PMSF, 0.1 mM AEBSF, 0.1 µM Aprotinin, 5 µM Bestatin, 2 µM E-64, 2 µM Leupeptin and 1 µM Pepstatin A). Cells were lysed by adding 0.5% Triton X-100 and incubating on ice for 10 min. Lysates were cleared by centrifugation (50,000 × g, 20 min, 4°C), and applied to 5 mL of Streptactin-XT resin (IBA Lifesciences, Göttingen, Germany) by gravity flow. The resin was washed with 50 mL of lysis buffer and protein was eluted with 25 mL of elution buffer (100 mM D-biotin in lysis buffer, pH 7.5). Eluted proteins were concentrated using 50K MWCO centrifugal filters (AS ONE Corporation #4-2669-05), flash frozen in liquid nitrogen, and stored at −80°C. Gel filtration was performed on a NGC chromatography system (Bio-Rad) using a BioSep-SEC-s4000, particle size 5 µm, pore size 500 Å, 7.8 mm ID x 600 mm column (Phenomenex) equilibrated in GF150 buffer (25 mM HEPES-KOH, 150 mM KCl, 2 mM MgCl2, pH 7.2). Fractions were analyzed by SDS-PAGE using the stain-free imaging with ChemiDoc (Bio-Rad). Peak fractions were collected and concentrated using an Amicon Ultra-0.5 (Merck Millipore). Protein concentration was determined by the absorbance of mScarlet at 569 nm using NanoDrop (Thermo Fisher Scientific). Proteins untagged with mScarlet were analyzed by BCA Assay (Nacalai Tesque) to determine the concentration. Proteins were flash frozen in liquid nitrogen and stored at −80°C.

#### Mass photometry

Proteins were diluted to 20-40 nM in GF150 buffer and analyzed using a Refeyn OneMP mass photometer (Refeyn Japan, Kobe, Japan) and Refeyn AcquireMP version 2.3 software, with default parameters set by Refeyn AcquireMP. Bovine serum albumin (BSA) was used as a control to determine the molecular weight. The results were subsequently analyzed using Refeyn DiscoverMP version 2.3, and graphs were prepared to show the distribution of molecular weight.

#### TIRF single-molecule motility assays

For preparing microtubules for TIRF assays, fresh pig brains were obtained from the Shibaura Slaughterhouse in Tokyo. Tubulin was purified from the brain as described. Tubulin was labeled with Biotin-PEG_2_-NHS ester (Tokyo Chemical Industry, Tokyo, Japan) and AZDye647 NHS ester (Fluoroprobes, Scottsdale, AZ, USA) as described (Chiba et al., 2022). To polymerize Taxol-stabilized microtubules labeled with biotin and AZDye647, 30 μM unlabeled tubulin, 1.5 μM biotin-labeled tubulin and 1.5 μM AZDye647-labeled tubulin were mixed in PEM buffer supplemented with 1 mM GTP and incubated for 15 min at 37°C. Then, an equal amount of PEM buffer supplemented with 40 μM taxol was added and further incubated for 30 min at 37°C. The solution was loaded on BRB80 supplemented with 30% sucrose and 10 μM taxol and ultracentrifuged at 20,000 g for 10 min at 25°C. The pellet containing polymerized microtubules was resuspended in PEM supplemented with 20 μM taxol and used for TIRF assays. Glass chambers were prepared by acid washing as previously described. Glass chambers were coated with PLL-PEG-biotin (SuSoS, Dübendorf, Switzerland). Polymerized microtubules were flowed into streptavidin adsorbed flow chambers and allowed to adhere for 5–10 min. Unbound microtubules were washed away using assay buffer (90 mM HEPES-KOH, 50 mM KCH_3_COO, 2 mM Mg(CH_3_COO)_2_, 1 mM EGTA, 10% glycerol, pH 7.4) supplemented with 0.1 mg/mL biotin–BSA, 0.2 mg/mL kappa-casein, 0.5% Pluronic F127, 2 mM ATP, and an oxygen scavenging system composed of PCA/PCD/Trolox. Purified motor protein was diluted to indicated concentrations in the assay buffer and flowed into the glass chamber. An ECLIPSE Ti2-E microscope equipped with a CFI Apochromat TIRF 100XC Oil objective lens (1.49 NA), an Andor iXion life 897 camera and a Ti2-LAPP illumination system (Nikon, Tokyo, Japan) was used to observe single molecule motility. NIS-Elements AR software ver. 5.2 (Nikon) was used to control the system.

#### Statistical analyses and graph preparation

Statistical analyses were performed using Graph Pad Prism version 10. Statistical methods are described in the figure legends. Graphs were also prepared using Graph Pad Prism version 10.

## Data availability

Plasmids used in this study have been deposited at Addgene. The data that support the findings of this study are available from the corresponding author upon request.

## Supporting information

This article contains supporting information, Figure S1, S2 and Table S1.

## Abbreviations and nomenclature

The abbreviations used are:

KIF: kinesin superfamily protein
MT: microtubule
CC: coiled-coil domain
TIRF: total internal reflection fluorescence
AEBSF: 4-benzenesulfonyl fluoride hydrochloride
PMSF: phenylmethylsulfonyl fluoride
ATP: adenosine triphosphate
MW: molecular weight

## Acknowledgments

The author is deeply grateful to Shinsuke Niwa and Richard J. McKenney for their invaluable discussions. I also thank Fumiko Takida for her assistance with this project, and Atsushi Nakagawa and Jiye Wang (Osaka University, Japan) for their support with mass photometry. I acknowledge the use of instrumentation provided by FRIS CoRE, a shared research facility at Tohoku University.

## Author contributions

K.C. designed research, performed experiments, analyzed data, and wrote the manuscript.

## Funding and additional information

KC was supported by MEXT (Ministry of Education, Culture, Sports, Science and Technology) Leading Initiative for Excellent Young Researchers (JPMXS0320200156), Naito foundation, Brain Science Foundation, Takeda Science Foundation (2024036450), Astellas Foundation for Research on Metabolic Disorders (2024A1048). This work was performed under the Collaborative Research Program of Institute for Protein Research, Osaka University, CR-23-02.

## Conflicts of Interest

The author declares no conflicts of interest.

### Declaration of generative AI and AI-assisted technologies in the manuscript preparation process

During the preparation of this manuscript, the author used ChatGPT (GPT-5.2) to check English grammar and improve the clarity of writing. After using this tool, the author carefully reviewed and edited the text and takes full responsibility for the final content of the publication.

