## Supplementary Materials (Figure S1, S2, Table S1) for "Activated kinesin-1 assembles into a dimer-of-dimers"

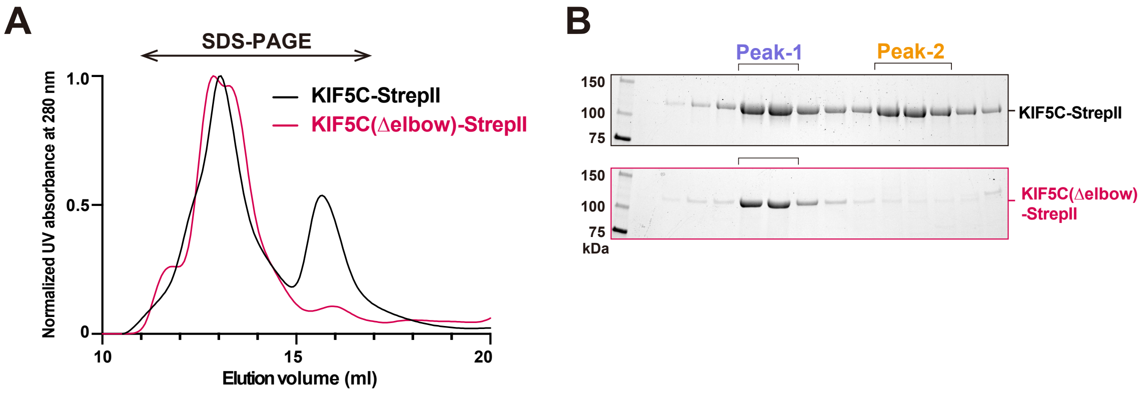


**Supplementary Figure S1. Removal of the fluorescent tag does not affect KIF5C tetrameric assembly.**

**(A)** Gel-filtration chromatograms of KIF5C-StrepII and KIF5C(Δelbow)-StrepII. Fractions indicated by double-headed arrow were analyzed by SDS–PAGE.

**(B)** SDS-PAGE of the fractions from (A). Peak positions are indicated by inverted U-shapes. Molecular weight standard (kDa) are shown on the left.

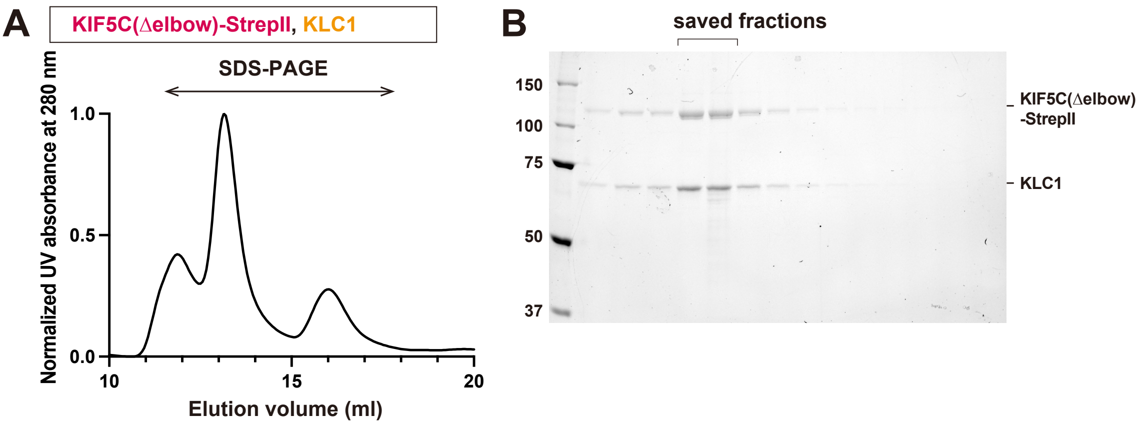


**Supplementary Figure S2. Association between KIF5C dimers is compatible with KLC binding.**

**(A)** Gel-filtration chromatogram of the KIF5C(Δelbow)-StrepII and KLC1 complex. Fractions indicated by the double-headed arrow were analyzed by SDS–PAGE.

**(B)** SDS-PAGE of gel filtration fractions from (A). Peak position is indicated by an inverted U-shape. Molecular weight standard (kDa) are shown on the left.

# Supplementary Table S1   Plasmid List

| Plasmid Name | Description | Comment | Label | Source |
|---|---|---|---|---|
| MOM657 | pACEBac1 Hs*KIF5A*-2xPPS-mScarlet-Strep-tag II | Figure 1 | KIF5A-mSca | Chiba et al., 2022 |
| MOM735 | pACEBac1 Hs*KIF5B*-2xPPS-mScarlet-Strep-tag II | Figure 1 | KIF5B-mSca | Chiba et al., 2022 |
| MOM658 | pACEBac1 Hs*KIF5C*-2xPPS-mScarlet-Strep-tag II | Figure 1 | KIF5C-mSca | Chiba et al., 2022 |
| pKC73 | pACEBac1 Hs*KIF5C*(ΔE675-R692)-2xPPS-mScarlet-Strep-tag II | Figure 3 | KIF5C(Δelbow)-mSca | This study |
| pKC97 | pACEBac1 Hs*KIF5C*-Strep-tag II | Figure 3, S1 | KIF5C-StrepII | This study |
| pKC98 | pACEBac1 Hs*KIF5C*(ΔE675-R692)-Strep-tag II | Figure 3, S1 | KIF5C(Δelbow)-StrepII | This study |
| pKC220 | pACEBac1 Hs*KIF5C*(1-914)-2xPPS-mScarlet-Strep-tag II | Figure 5 | KIF5C(Δtail)-mSca | This study |
| pKC102 | pACEBac1 Hs*KIF5C*(1-914,ΔE675-R692)-2xPPS-mScarlet-Strep-tag II | Figure 5 | KIF5C(ΔtailΔelbow)-mSca | This study |
| pKC168 | pACEBac1/pIDS<br>pACEBac1 Hs*KIF5C*(ΔE675-R692)-Strep-tag II<br>pIDS His-FLAG-Hs*KLC1* | Figure S2 | KIF5C(Δelbow)-StrepII, KLC1 | This study |
